# Long-term phenotypic and genomic instability of an industrial ethanol-producing and C5-utilizing *Saccharomyces cerevisiae* strain

**DOI:** 10.1101/2023.09.04.555844

**Authors:** Maëlle Duperray, Mathéo Delvenne, Jean-Marie François, Frank Delvigne, Jean-Pascal Capp

**Affiliations:** Toulouse Biotechnology Institute, INSA/University of Toulouse, CNRS, INRAE, Toulouse, France; TERRA research and teaching centre, Microbial Processes and Interactions (MiPI), Gembloux Agro- Bio Tech, University of Liège, Gembloux, Belgium; Toulouse White Biotechnology, INSA, INRAE, CNRS, Toulouse, France

**Keywords:** phenotypic heterogeneity, genetic stability, industrial yeast strain, homologous recombination, xylose, arabinose, Ethanol Red

## Abstract

The production of bio-based compounds can be impacted by stochastic cellular mechanisms acting at both the genetic and non-genetic levels, leading to instabilities in the performance of the producer organisms. Here, we investigated the long-term genomic and phenotypic stability of an ethanol-producing *Saccharomyces cerevisiae* strain engineered by chromosomal integration of transgenes allowing xylose and arabinose utilization. First, along a series of batch cultures in bioreactor to reach around 100 generations, we observed that, after a period of phenotypic stability, the strain exhibited instabilities in terms of xylose and arabinose consumption. Long-read sequencing did not reveal major genomic modifications at the population level that could explain such fluctuations. However, we isolated clones that have partly or fully lost the ability to use arabinose or xylose, due to copy number variations of the integrated transgenes arising most probably from homologous recombination (HR) events. Based on the cultivation of subpopulations sorted depending on their expression level of *RAD52*, a gene whose expression is known to be proportional to the HR rate, we did not detect differences in genomic and phenotypic stability in the subpopulations. Thus, this work reveals both phenotypic and genomic variations in an industrial yeast strain that could in the long-term lead to the loss of its production performance.

## Introduction

Industrial strain stability is a central issue in modern biotechnology. In the framework of synthetic biology for microbial production, reduced production titer and yield can occur due to spontaneous variants that will escape the metabolic burden caused by expression of the synthetic pathway and concurrent potential toxicity associated with accumulation of non-natural intermediates and end-products [1, 2]. Genetic instability has been the main focus of previous studies to provide mechanistic explanation of strain instability during bioprocessing production. In *Escherichia coli*, it was reported that the rise of frequencies of mobile element insertions in chromosomally-inserted transgenes is the main factor that interrupts production in mevalonic acid-producing cells [1]. Similarly, it has been shown that the metabolic stress induced by L-cysteine production in *E. coli* triggers insertion sequence (IS) transposition, resulting in structural genetic rearrangements in production plasmids that cause L-cysteine production capacities to drop by up to 65–85% within 60 generations [3]. Consequently, selective deletion of IS and their corresponding transposases could prove useful for stabilizing industrially production strains [3]. In *Saccharomyces cerevisiae*, homologous recombination (HR) has been recently reported as an important genetic process leading to the loss of production capacity [4], as illustrated in the case of vanillin-β-glucoside [5]. The HR-induced instability of producer strains is partly explained by the fact that cassettes carrying the genes of the production pathway integrated into multicopy in the genome and carrying identical promoter/terminator are easily excised under industrial production conditions, which are particularly highly constraining [2].

In addition to genetic instability, the last decade has shed light on the origins of low-performing variants, showing the non-genetic sources of strain instability assigned to cell-to-cell heterogeneity in biotechnological production. Notably, metabolic heterogeneity with clonal microbial cells varying in metabolite levels [6] and metabolic fluxes has been highlighted [7] with causal factors ranging from intracellular molecular noise, especially stochastic gene expression [8], to environmental factors [9]. Genetic and non-genetic heterogeneity are generally considered independently as origins of strain instability. However, there is an interplay between genetic, epigenetic and gene expression variability [10] that complicates the study and the control of such instability. Especially, noise in the expression of genes involved in DNA repair and recombination [11] can result in cell-to-cell heterogeneity both in bacteria [12] and yeast [13]. For instance, it was found that spontaneous HR rates could vary up to 10-fold in a *S. cerevisiae* clonal population depending on the level of expression of genes affecting HR activity either directly (such as *RAD52* involved in the early steps of HR pathways) or indirectly (such as *RAD27* involved in Okasaki fragments maturation) [13]. This HR-dependent cell-to-cell heterogeneity may thus favor the emergence of adaptive behavior to environmental conditions, as it can trigger the occurrence of low-performing variants which might be responsible for progressive genetic drift of the population.

In this report, we addressed the long-term genomic and phenotypic stability of an ethanol-producing strain derivative from the Ethanol Red^®^ strain engineered to co-ferment C5 sugars (xylose and arabinose) with glucose. We found that the rate of xylose and arabinose consumption started to fluctuate after around 50 generations without any major genomic modifications detected at the population level. However, we isolated clones exhibiting partial or full loss of ability to use arabinose and/or xylose due to transgene copy number variations (CNV). Contrary to expectations, subpopulations enriched in either high or low *RAD52* expression, whose variation was reported to trigger heterogeneity in HR rate [13] did not exhibit differences neither in C5 consumption nor in genomic content. This indicates that other factors than heterogeneity in *RAD52* expression are responsible for these CNV, which nonetheless could lead to the loss of the fermentation capacity of these sugars in the long- term.

## Material & Methods

### Strains, plasmids and media

All experiments were performed using the *Saccharomyces cerevisiae* HDY.GUF12 strain [14, 15] (Lesaffre France). The HDY.GUF12 strain containing the *RAD52-YFP-tdTomato* fusion was obtained using a CRISPR-Cas9 strategy [16]. The high copy pCas9-amdSYM plasmid (derived from pML107 [17]), which constitutively expresses the gene encoding the Cas9 endonuclease and carrying a guide RNA (gRNA) expression cassette, was used to transform the HDY.GUF12 strain. A healing fragment containing *YFP-tdTomato-SpHis5* and homologies to *RAD52* was amplified from the JA0241 strain [13] (primers GATCCCAAATACCAGGCACA and CATAGTTCAATTGCGTGACATC) and used to repair the double-strand break introduced by the Cas9 endonuclease. The target sequence of the gRNA was identified using the online software CRISPR-direct (https://crispr.dbcls.jp), and located in the *RAD52* 3’UTR region. Ligation of the gRNA expression cassette sequence into the pCas9-amdSYM plasmid was made using a T4 DNA ligase (NEB). Yeast cells were transformed using the yeast transformation procedure described in [14] and selected on YNB Acetamide plates (1.7 g/L yeast nitrogen base without amino acids and nitrogen (Euromedex), 6.6 g/L K2SO4, 0.6 g/L acetamide and 20 g/L D-glucose) as the amdSYM cassette confers the ability to use acetamide as the sole nitrogen source [18]. Strain and plasmid constructions were verified by PCR amplification and sequencing. A synthetic mineral yeast medium adapted from the Verduyn recipe [19] was used for the serial cultivations in microplates. This medium, named SHD2, contained 5-times more trace elements than the original recipe, 0.1 M potassium hydrogen phthalate, 16 g/L D-glucose, 13 g/L D-xylose and 11 g/L L-arabinose and adjusted at pH 5.0. Rich YPD medium contained 20 g/L D-glucose, 20 g/L bactopeptone and 10 g/L yeast extract. Minimal YNB medium contained 1.7 g/L yeast nitrogen base without amino acids and nitrogen (Euromedex) and 5 g/L ammonium sulphate. YNB was supplemented with appropriate sugars, as indicated in the following sections.

### Batch fermentation

Cultivations were performed in 750 mL 2X YNB medium supplemented with 60 g/L D-glucose, 26 g/L D-xylose and 22 g/L L-arabinose in a 1 L bioreactor (Solaris Biotech Solutions). Agitation and temperature were set at 400 rpm and 30 °C respectively and pH was maintained at 5.5. Airflow was maintained at 100 mL/min during the first 18 h (until all glucose was consumed), before assigned anaerobic conditions (N2 injection). Bioreactors were connected to an online gas analyzer (Prima PRO, Thermo Scientific) to dose ethanol in gas. Cells were inoculated in the first batch at OD = 0.1 from overnight pre-culture of the WT HDY.GUF12 strain and cultivated for 48 h before being inoculated in the next batch at the same initial OD. Serial batch cultivations were performed in duplicate allowing to reach 90 (15 successive batch) or 96 (16 successive batch) (TableS1). Samples were regularly collected all along cultivations to perform OD measurements and further metabolite quantifications. At the end of each batch, cells were also stocked in glycerol solution and kept at -80 °C.

### Successive cultivation in microplates and cell sorting

To initiate the successive cultures in microplates, cell suspensions of each strain (WT and *RAD52-YFP-tdTomato* strains) obtained from overnight pre-cultures in SHD2 medium, were inoculated in a 48-well FlowerPlate (800 µ L at OD600 = 0.1). Cultures were performed in the Biolector Pro (Beckman Coulter GmbH, Germany) at 30 °C in SHD2 medium, under propagation conditions for 24 h at a shaking frequency of 1000 min^-1^ to ensure a high oxygen transfer rate. This was followed by cultivation under production conditions for the last 48 h at a shaking frequency of 800 min^-1^ after the addition of 800 µ L of fresh medium to decrease the oxygen transfer rate. After this initial culture cycle (C0), OD (600 nm) was measured, cells were harvested and the supernatant was kept at -20 °C for further HPLC analysis while the pellet was resuspended in PBS and kept at -20 °C before cell sorting. At this stage, cells were also stocked in glycerol solution and kept at -80 °C. The cells were sorted using a FACS Aria III (BD biosciences, Belgium) based on a custom mode (0-16-0) with an 85 µm nozzle. A first gate was applied to select cells based on YFP (488 nm laser, 530/30 filter) and tdTomato (561 nm laser, 582/15 filter) fluorescence signals. Based on the FSC-Area vs. FSC-Height (561 nm laser) plot, single cells with similar cell size and granularity were then selected. Finally, based on the histogram of the YFP-tdTomato fluorescence (561 nm laser, 582/15 filter), single cells with the 10% highest (renamed Rad52-high) and the 10% lowest (renamed Rad52-low) Rad52-YFP-tdTomato fluorescence levels were sorted simultaneously and recovered in PBS. The WT strain was sorted without this final gating (100% WT single cells). After cell sorting, recovered cells (approximately 2.10^5^ cells for each strain (Table S2 for details)) were harvested (10,000 g during 30 min) and resuspended in 800 µ L of fresh medium to be inoculated in a 48-well FlowerPlate. These cells were cultivated, sampled and sorted as the C0 culture cycle, except in the following steps: propagation condition lasted 48 h; only the high-Rad52 cells were recovered from the cultures of Rad52-high, while only the low-Rad52 cells were recovered from the cultures of Rad52-low, thus enriching the cultures with high- or low-Rad52 cells respectively. A total of nine culture cycles have been performed for each condition (Rad52-low, Rad52-high and WT strain) to reach more than 90 generations.

### Replica tests and growth analysis by spot assays

About 1500 cells from an overnight culture were spread on YNB plates containing 20 g/L D-glucose, 10 g/L D-xylose and 10 g/L L-arabinose. The clones from each plate were then transferred by replica plating on YNB plate containing 20 g/L D-glucose, 20 g/L D-xylose or 20 g/L L-arabinose. Clones exhibiting a growth difference on xylose and/or arabinose medium to that on glucose medium were selected and further analyzed by spot assays. They were grown overnight on YPD plates and resuspended in sterile MilliQ water at 2.10^7^ cells/mL before being submitted to 1/10 serial dilutions. Drops (5*µ*L) of each dilution were spotted onto freshly prepared YNB plates containing 20 g/L D-glucose, 20 g/L D-xylose or 20 g/L L-arabinose and were incubated at 30 °C.

### Metabolite analysis

Supernatants from cultures were syringe filtered (0.2 µm) before analysis using HPLC. Samples from the successive cultivations in microplate were analyzed using an HPLC Agilent 1200 Series (Agilent) equipped with an Aminex HPX-87H column coupled with a pre-column (Bio-Rad). The other samples were analyzed using either Vanquish or Ultimate 3000 HPLC system (Thermo Scientific) equipped with a REZEX ROA-Organic Acid H+ column coupled with a pre-column (Phenomenex). For all experiments, the eluent solution was 5 mM H2SO4, running at 0.5 mL/min for 40 min at 50 °C. Compounds were detected by an RI detector and quantified from standard curves using the Chromeleon software.

### Real-time PCR assays

Genomic DNA was extracted using the MasterPure Yeast DNA Purification kit (LGC Biosearch Technologies) and quantified by NanoDrop (Thermo Scientific). The copy number of the five genes of interest were determined using the MyIQ real-time PCR system from Bio-Rad. The reaction mix (25 *µ*L final volume) contained 12.5 *µ*L of iQ SYBR Green Supermix buffer (Bio-Rad), 3 *µ*L of each primer (Table S3) at a final concentration of 250 nM and 10 ng of DNA. The thermocycling program consisted of one hold at 95 °C for 5 min, followed by 40 cycles of 10 s at 95 °C and 45 s at 56 °C. To correlate the copy number of each gene to the Ct value, calibration curves were performed with each couple of primers on serial dilutions of DNA from the WT strain (FigS5). Genome copy number was first determined using the *ACT1* gene as a reference, allowing us to calculate the copy number of each gene of interest. The resulting data were then normalized to the WT initial strain condition for which copy number of genes is known.

### Genome sequencing

Cells were cultivated in YPD medium directly from glycerol stock to obtain 10^9^ cells.

Genomic DNA was extracted using the Blood & Cell Culture DNA Midi Kit (Qiagen) according the manufacturer’s protocol adapted for yeast cells with following modifications: (i) lyticase digestion extended to 2 h; (ii) proteinase K digestion extended to 3 h; (iii) cellular debris pelleted by centrifugation at 5,000 *g* for 20 min with supernatant filtered (0.2 µm) before applying to the equilibrated genomic-tip; (iv) DNA precipitated in isopropanol was pelleted by centrifugation at 5,000 g for 20 min, washed two-times in cold ethanol 70%, dried at 37°C for 30 min and dissolved overnight in EB buffer (Qiagen). DNA was quantified by NanoDrop (Thermo Scientific) and its quality control was validated on Qubit (Thermo Scientific) and Fragment Analyzer (Agilent).

Single-molecule real-time long reads sequencing was performed at Gentyane Sequencing Platform (Clermont-Ferrand, France) with a PacBio Sequel II Sequencer (Pacific Biosciences, Menlo Park, CA, USA). The SMRTBell library was prepared using an SMRTbell prep kit 3.0, following the procedure and checklist -preparing whole genome and metagenome libraries protocol. Genomic DNA (1 µg) of each strain was sheared using g-tubes (Covaris, England) generating DNA fragments of approximately 10 kb. Sheared genomic DNA was carried into the enzymatic reactions to remove the single-strand overhangs and to repair any damage that may be present on the DNA backbone. An A-tailing reaction followed by the barcoded overhang adapter ligation was conducted to generate the SMRT Bell templates. After nuclease treatment, the sample was then size-selected for fragments above 5kb with 35% AMPure PB Beads and equimolar multiplied to obtain the final libraries around 10 kb. A ready-to-sequence SMRTBell Polymerase Complex was created using a Binding Kit 3.2 (PacBio) and the Sequel II primer 3.2. The PacBio Sequel instrument was programmed to load a 90 pM library and sequenced in CCS mode on a PacBio SMRTcell 8M, with the Sequencing Plate 2.0 (Pacific Biosciences), 2 hours of pre-extension time and acquiring one movie of 15 hours per SMRTcell.

### Bioinformatics analysis

The CCS tool (v6.3.0) from the smrttools suite (11.0.0.146107), with default parameters, was used to generate the corrected CCS data (20.03 Gb, mean 8,880 bp) from raw data. These CCS were demultiplexed per sample using lima (2.5.1). Assembly with hifiasm (0.16.1-r375) was performed to obtain the two haplotypes of diploid strains. Each contig was aligned against the reference strain S288C, using nucmer (3.1). Thus, a chromosome name could be assigned to each contig, based on the percentage identity between S288C and the hifiasm assemblies. The pbmm2 tool (2.8.1) was used to align the pools of reads against the diploid reference assembly (*RAD52-YFP-tdTomato* strain). The pbsv pipeline (2.8.1) was used to call structural variants with a minimal size set at 2 nucleotides. The various vcf calling files were collected and analysed using VCFtools (0.1.16). The various SVs found were visualised and confirmed using IGV.

## Results

### Investigation of the long-term phenotypic and genomic stability of an industrial strain

To assess the phenotypic and genomic stability of an industrially relevant yeast strain in the context of bioproduction, we choose an ethanol-producing strain, derivative from the diploid Ethanol Red^®^ strain, one of the most used yeast strains for the first-generation bioethanol production. This strain named HDY.GUF12 has been previously engineered to co-ferment C5 sugars (xylose and arabinose) with glucose, by chromosomal transgene insertions and evolutionary engineering experiments [14, 20] (FigS1). HDY.GUF12 strain was cultivated in duplicate (Dupl.1 and Dupl.2) for 16 successive batch bioreactors to reach more than 90 generations (Fig1A, Table S1), mimicking the typical succession of generations considered for industrial fermentation processes [2]. Each batch lasted 48 hours with an aerobic phase that lasted about 18 hours to promote growth, followed, at the beginning of the stationary phase, by an anaerobic phase during which only ethanolic fermentation took place.

**Figure 1:**
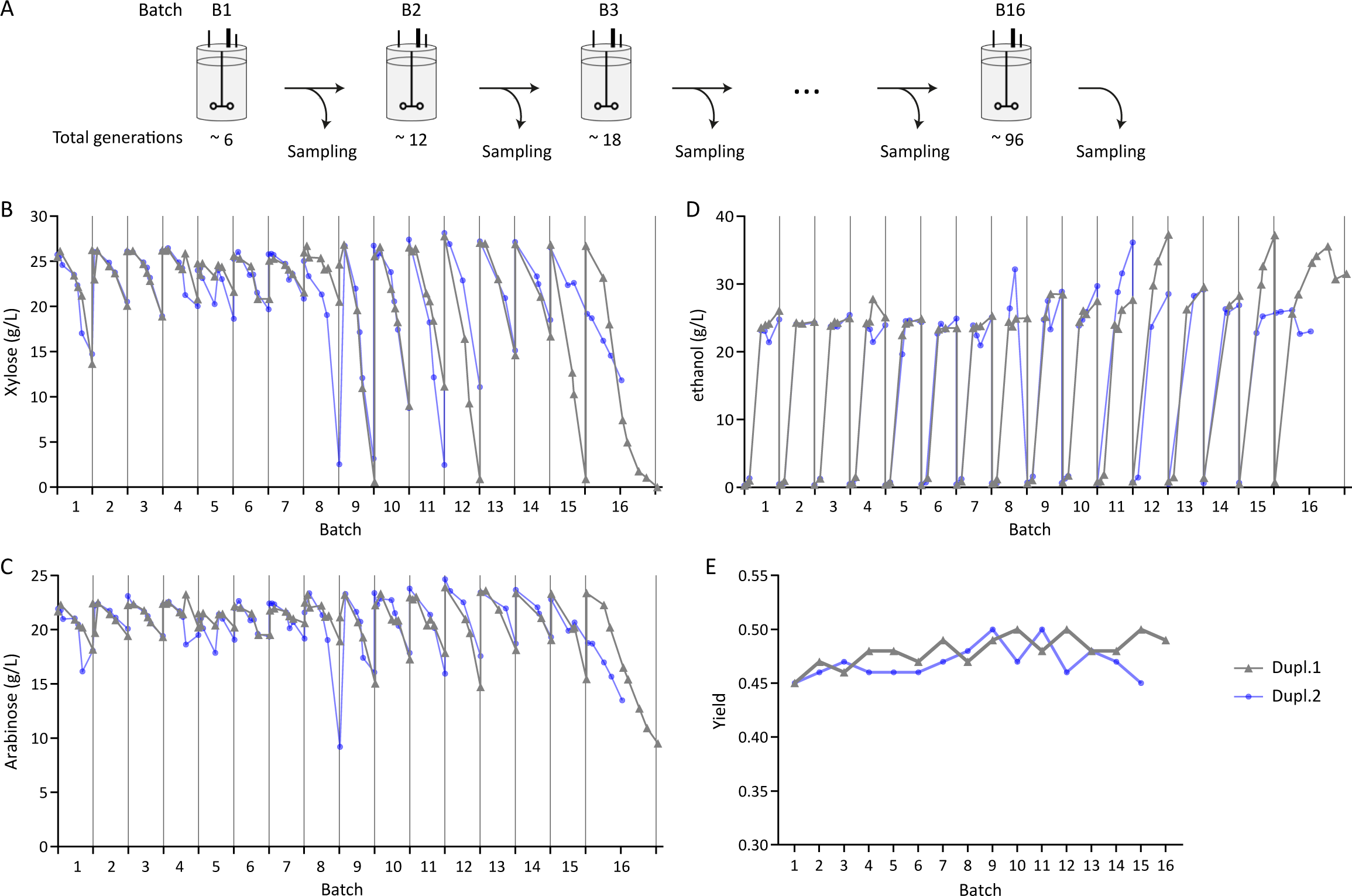
Heterogeneity in xylose and arabinose consumption and ethanol production in successive batch cultures. A: Serial cultivations performed in batch reactor to reach about 96 generations. Samples from two parallel cultivations (Dupl.1 and Dupl.2) were regularly collected for subsequent analyses. Extracellular xylose (B), arabinose (C) and ethanol (D) quantification along cultivations. E: Ethanol yield estimated from the last measures of each batch and calculated by dividing the total number of carbon produced by the total number of carbon consumed from glucose, xylose and arabinose.

While glucose was always totally consumed at the beginning of the stationary phase (data not shown), the xylose and arabinose consumption fluctuated from one batch to the other, from moderate to total consumption (Fig1B-C). These fluctuations were far more visible in the second half of the batches (8 to 16) and followed the same trend for xylose and arabinose i.e., the more xylose was consumed during a batch, the more arabinose was also consumed.

Moreover, the final OD was similar between batches and duplicates (Table S1), meaning that xylose and arabinose were not significantly used for biomass formation. Instead, they were mainly used for ethanol production as the ethanol titer is correlated to C5 sugars consumption (Fig1D). In addition, the yield of ethanol was measured in the range of 45-50% of sugars consumed all over the successive batch (Fig1E). It was also verified that the ethanol tolerance of cell population taken along the successive batches was comparable (FigS2), indicating that observed fluctuations in C5 consumption along successive batch was not associated with toxic effect due to high ethanol production but rather due to differences in the ability to consume xylose and arabinose.

To determine if these differences can be explained by genomic variations, we performed whole genome long-read sequencing to analyze the global genomic stability at the population level. For both, sequencing was performed on samples coming from four representative batch cultures, chosen for their different cultivation time and phenotypes (B5, B9, B13 and B16 for Dupl.1; B5, B9, B12 and B15 for Dupl.2). No genomic differences were identified between the tested samples, indicating a strong stability of the strain at the population level despite the phenotypic variability we observed.

### The bulk population contains C5 low-consuming cells that exhibit copy number variation of arabinose-associated transgenes

The peculiarity of C5 sugars consumption over the successive batches prompted us to scrutinize the possibility that some clones in the clonal population had possibly lost their capacity to consume these sugars. To this end, a screen was performed to isolate clones that have lost, at least in part, the ability to metabolize xylose and/or arabinose (FigS3A). Cells from the eight previously selected batches (B5, B9, B13 and B16 for Dupl.1; B5, B9, B12 and B15 for Dupl.2) were first plated on a medium containing glucose, xylose and arabinose and replica-plated on media containing only glucose, xylose or arabinose as the sole carbon source. Clones exhibiting a differential growth on xylose and/or arabinose as compared to glucose were further analyzed by spotting serial cell dilutions (FigS3B). This allowed isolating several clones that indeed exhibited a growth defect on at least one of the C5 sugars with a proportion being 1.49% at the maximal of the total spread cells per batch (FigS3C).

However, there was no correlation between the percentage of these clones in a given batch and the corresponding C5 sugars consumption, probably because the actual number of low-consuming clones was underestimated in such screening. From a total of 30 clones isolated, two-thirds were affected in their ability to grow on xylose and all of them were affected in their ability to grow on arabinose (FigS3B and D). We chose nine of them exhibiting different growth phenotypes and isolated from three different batch cultures for further analysis. The loss of their ability to metabolize xylose and/or arabinose on plate was first confirmed (Fig2A). We then quantified sugars consumption and ethanol production after 24 hours in medium containing a mix of glucose, xylose and arabinose (Fig2B). While there was no difference in consumption between the initial strain and the bulk populations from the different batches, all the nine selected clones exhibited a lower consumption of either xylose, arabinose or both (Fig2B), in accordance with the spotting results (Fig2A). Ethanol production was also altered, in correlation with C5 sugars consumption (FigS4). This suggests that only their ability to metabolize xylose and/or arabinose was affected while the yield of these clones was comparable to the initial strain.

**Figure 2:**
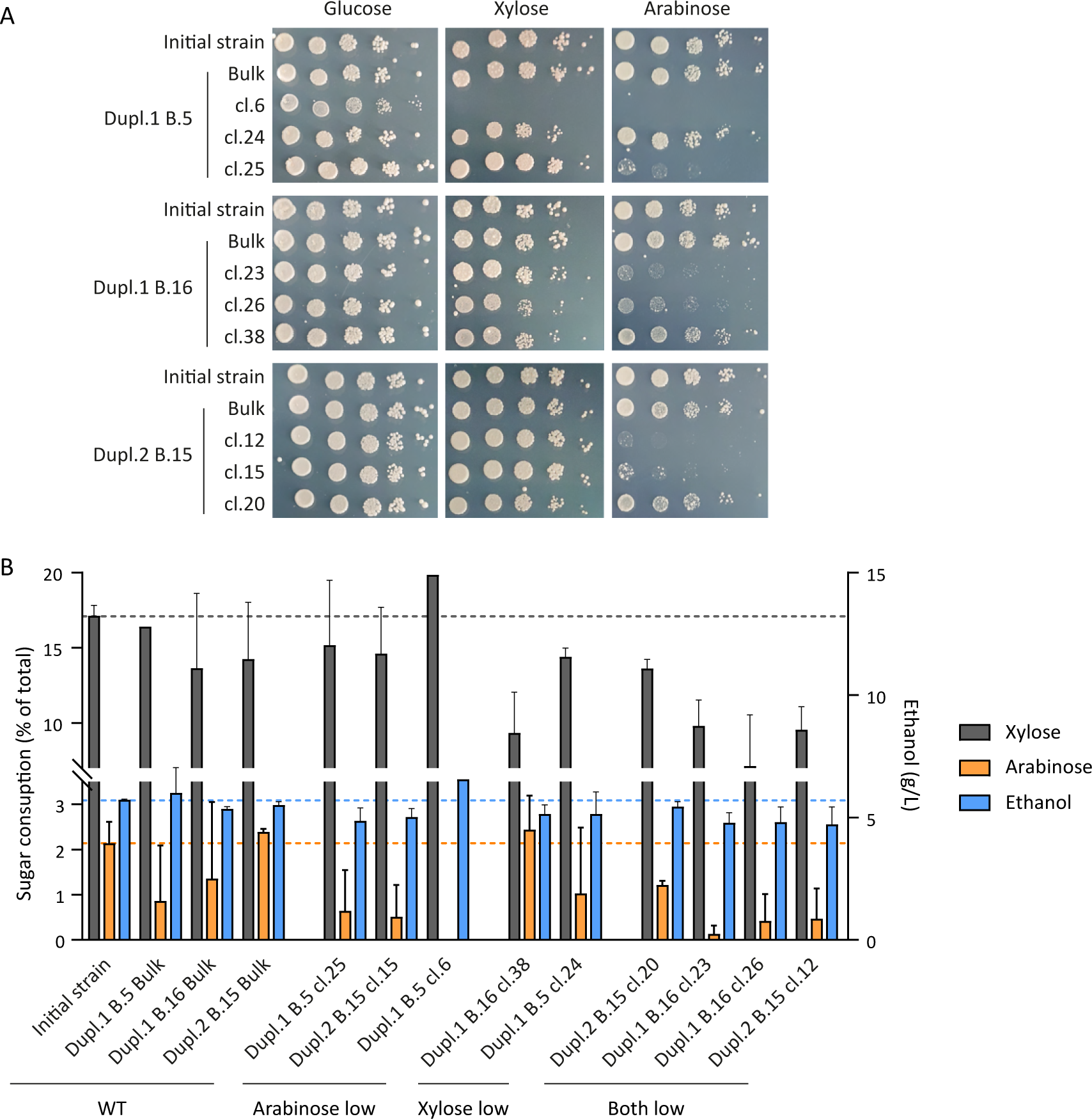
Xylose and arabinose consumption of selected clones isolated from batch cultures. A: Spotting assays of serial dilutions of cell suspension on YNB plates containing the indicated sugar at a final concentration of 20g/L. B: Extracellular xylose, arabinose and ethanol quantification after 24 h on YNB medium containing 16 g/L D-glucose, 13 g/L D-xylose and 11 g/L L-arabinose (n=2).

We then investigated whether these low-consuming phenotypes of the clones were associated with the loss of transgenes involved in C5 utilization. As most of the clones exhibited an arabinose low-consuming phenotype, we focused on arabinose-associated transgenes. The *araA*, *araB*, *araD* and *araT* copy numbers were determined by qPCR based on calibration curves (FigS5) and using *ACT1* as a reference for normalization. While the copy number of the tested transgenes was the same between the bulk populations from the batch cultures and the initial strain, most of the tested clones exhibited lower *araB*, *araA* and *araD* copy numbers (Fig3). In particular, each clone displayed a CNV for at least one gene among these three genes with the most affected being *araB,* encoding L-ribulokinase, followed by *araA,* encoding the arabinose isomerase giving rise to L-ribulose from L-arabinose, whereas the copy number of araT gene that encodes the transporter showed no variation. Previous findings shown that the sugar consumption phenotype of HDY.GUF12 strain was correlated with *araA* (and *xylA*) copy number [14], pointing out the importance of *araA* in this consumption. Altogether, these data indicated that a decrease in the arabinose-associated transgenes copy number could account explain the low-consuming phenotype in the isolated clones. Thus, rare cells in the population displayed genome modification, at least in transgene copy number, that affected their ability to consume C5 sugars.

**Figure 3:**
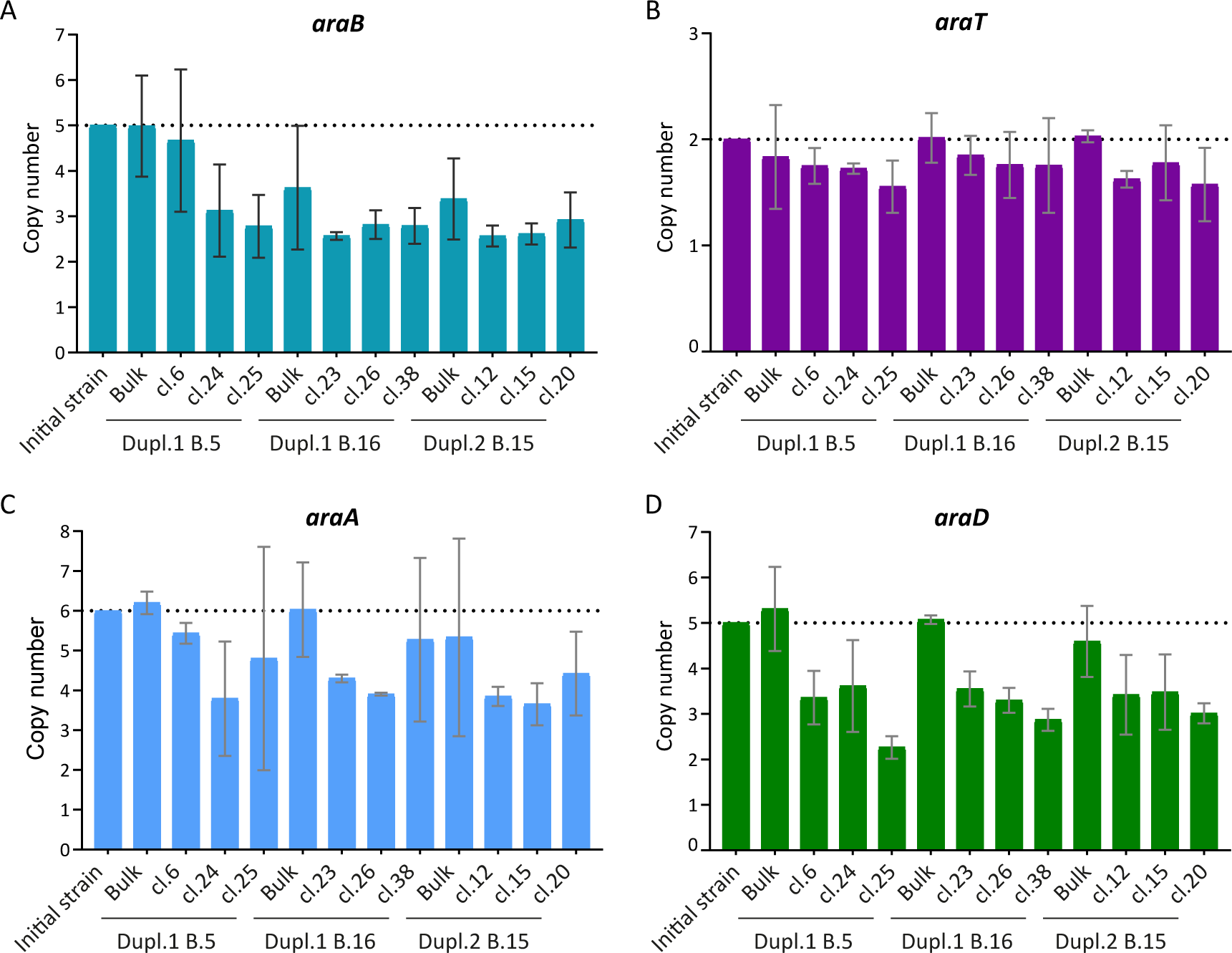
Copy number of the transgenes involved in arabinose consumption in selected clones isolated from batch cultures. Copy number of *araB* (A), *araT* (B), *araA* (C) and *araD* (D) were determined by qPCR. *ACT1* served as a reference gene and data were normalized to the WT initial strain condition (n=3).

### Comparison of C5 consumption between subpopulations enriched in cells expressing high or low Rad52 levels

In yeast, CNVs are mostly due to HR events. These events are more frequent in diploid strains and when transgenes are inserted in multiple copies such as in HDY.GUF12 (FigS1). . It has been previously shown that the rate of spontaneous HR in an *S. cerevisiae* clonal population positively correlates with the expression level of genes involved in HR activity such as *RAD52* [13]. To investigate whether the subpopulations expressing more *RAD52* accumulate more genomic modifications, we performed successive cell sorting consisting of regularly enriching the population either in high- or low-Rad52 expressing cells. Cells were successively cultivated in microplate and sorted based on the expression of Rad52 thanks to the integration of the *YFP-tdTomato* reporter cassette in C-ter at the *RAD52* original locus. Nine cycles of cultivations were performed with successive sorting of the 10% Rad52-highest expressing cells (Rad52-high) and the 10% Rad52-lowest expressing cells (Rad52-low) in parallel, while the untagged strain (WT) was cultivated without cell sorting (Fig4A and FigS6). This experiment allowed us to reach more than 90 generations for each condition (TableS2) with significant enrichment in high- and low-Rad52 cells respectively, as shown by the fluorescence profile of the populations (FigS7). The dosage of C5 sugars was performed at the end of each cycle, and no difference was detected between the three populations (Fig4B and 4C), indicating that the enrichment in Rad52-high cells did not lead to visible phenotypic effects in terms of sugars consumption that would be due to higher genetic instability. Of note, the final OD for Rad52-high populations progressively decreased compared to the Rad52-low and the non-sorted populations (Fig4D and TableS2).

**Figure 4:**
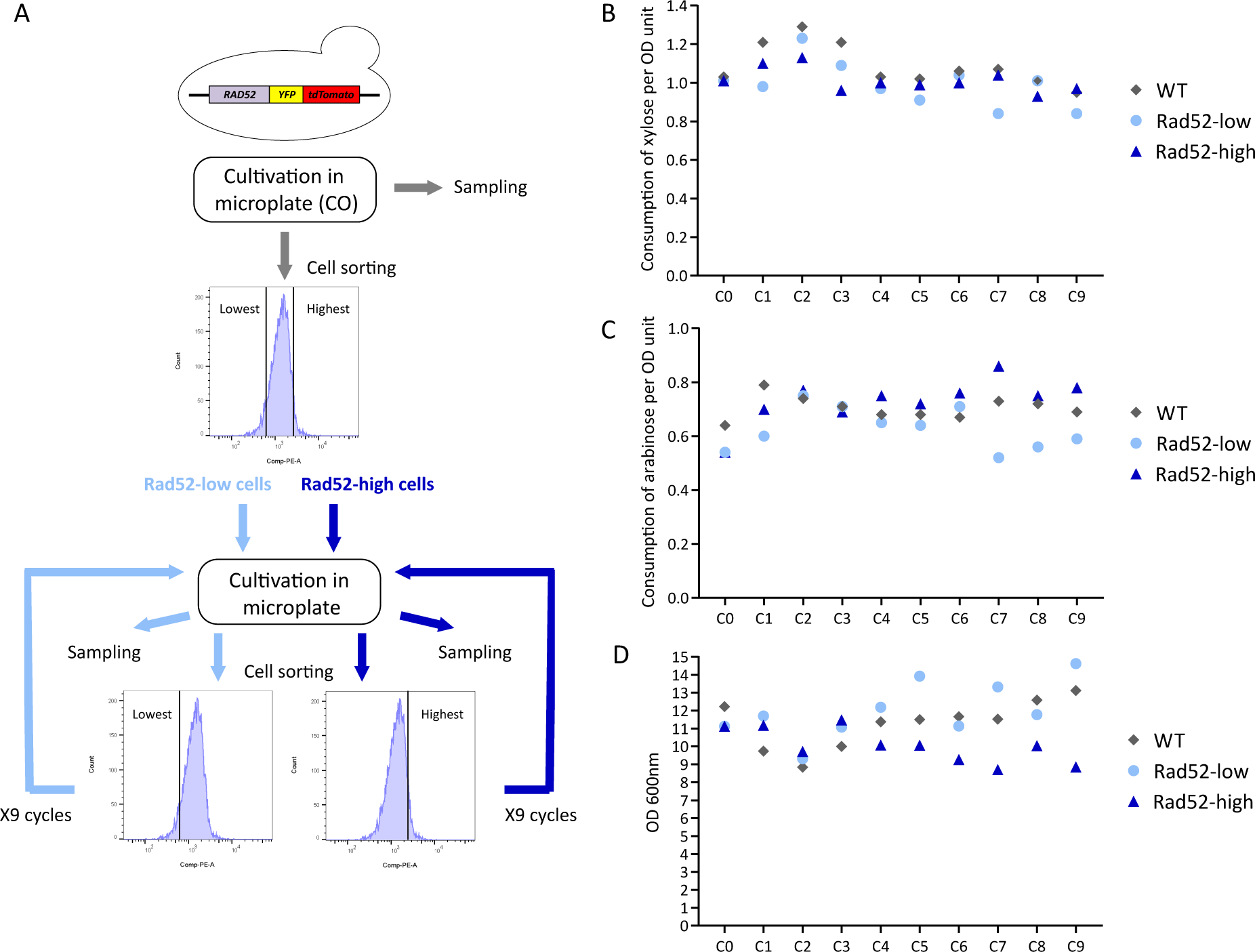
Serial cultivations of subpopulations enriched in either high- or low-Rad52 expressing cells. A: Experimental workflow of the successive cell sorting based on the Rad52 expression level and cultivations in microplate experiment. Samples were collected at the end of each cycle for subsequent analysis. Extracellular xylose (B), arabinose (C) and OD (D) quantification from samples collected at the end of each cycle.

We then wondered if this enrichment led to the appearance of genomic modifications in the Rad52-high population during cultivation despite the absence of phenotypic effects on sugars consumption. We performed long-read sequencing on samples from the C0 and C9 cycles for the WT strain and from the C0, C3, C7 and C9 cycles for the Rad52-high and Rad52-low cultures. No difference was observed for the transgenes, in accordance with the stability of C5 sugars consumption. Nevertheless, three loss of heterozygosity (LOH) in the WT strain from the C9 culture compared to the C0 culture was found, one in the intergenic tandem repeats region between *YPR015C* and *TIF6*, one in the promoter of *CMR3*, which is a putative zinc-finger protein, and the last one on the telomere region on the right arm of chromosome X (TableS4). These genetic modifications probably occurred through HR and invaded the population of this strain. Moreover, we identified two LOH regarding transposon sequences in the Rad52-low population from the C3, C7, and C9 cultures. No genomic difference was observed between the populations from Rad52-high cultures. Thus we did not detect any enrichment of HR-linked genomic modifications in the Rad52-high cells, suggesting that other factors than heterogeneity in *RAD52* expression are responsible for the CNVs we observed in the WT strain.

## Discussion

The genetic and phenotypic stability of microorganisms designed for the production of value-added products from carbonaceous biomass is a crucial issue in industrial biotechnology and one that is the subject of a growing body of work. Initial work on this subject focused on the stability of production systems in bacteria, where it was clearly shown that the use of plasmids carrying these production pathways is not advisable (see for instance [3]). Work on biomolecules-producing yeasts is less advanced, with a few works also revealing the problem of the stability of producing strains when the metabolic system is plasmid-based expressed [4, 5]. In this study, we set out to address the issue of genetic and phenotypic stability using a yeast designed to produce bioethanol from lignocellulose sugars by the genomic insertion of several copies of C5 sugars assimilation genes [14, 15]. This question was assessed by successive cultivation in batches to reach up to 96 generations, mimicking the generations occurring in industrial fermentation processes.

The strain harbored a quite stable but moderate phenotype in terms of C5 consumption along the first half of successive cultivations performed in batch reactors, contrary to previous work where C5 consumption was complete during short cultivation of this strain [14]. We observed that the consumption phenotype started to fluctuate after around 50 generations, from limited to full consumption with a highly non-reproducible phenotype from one batch to the next. Thus, as soon as the strain switched to higher consumption, it appeared that its phenotypes became highly variable. These results suggest a non-genetic origin from these fluctuations because each batch is likely too short to generate selection or counter-selection of genetic variants that could explain such variations in terms of sugars consumption. This hypothesis has been confirmed by long-read sequencing that did not reveal genomic differences along the batch cultivations at the population level. These sequencing results also confirmed the high genetic stability expected for an industrial strain used worldwide for ethanol production, but this does not prevent phenotypic fluctuations such as the ones observed in this work. No difference in yield was observed i.e., only the ability to transport or metabolize C5 sugars fluctuated and not the ability to produce ethanol. Beyond the transgenes integrated, many endogenous genes involved in C5 sugars consumption [21] might be the targets of epigenetic events that could modulate their expression level and explain the observed phenotypic variations. Comparative transcriptomics analysis would help decipher the non-genetic origin of these variations.

Despite the absence of genomic modifications detected at the bulk level after 96 generations, we were able to isolate clones with rare genetic variants harboring a reduction of copy number of transgenes, impacting the assimilation of these C5 sugars. While these variants are too rare to impact the sugars consumption of the bulk population and to be revealed by sequencing, it cannot be excluded that in a longer term, the abundance of these variants would increase, especially because under anaerobiosis, energetic provision from C5 sugars is 12.5 % less per mol sugar than from glucose. Cells losing the capacity to assimilate C5 sugars should have fitness advantage over those keeping this ability. Thus, a basal low level of genetic instability exists and might, depending on the environmental conditions, produce a genetic drift of the population, especially at large scale in industrial bioreactors where more challenging conditions would appear.

Cells isolated for high or low Rad52 expression did not exhibit differences in C5 consumption and ethanol yield, showing that even weak, the CNVs could not be ascribed to heterogeneity in Rad52-based HR events. However, we might have not cultivated the populations long enough or in sufficiently challenging conditions to allow enrichment of HR-related genomic events. Especially, we cannot exclude that the HR rate was not high enough to generate many genomic alterations in the duration of the experiment (only three LOH events were detected in the non-sorted populations). Also, the HR rate might not be different between the populations sorted for their Rad52 expression level in this industrial strain, even if it has been shown in the BY laboratory strain [13]. The only difference observed was the progressive decrease in final OD for the Rad52-high population along the cycles. This decrease might be due to the higher expression of the YFP-tdTomato tag, higher Rad52 levels or indirect consequences of this higher expression (other than genetic because sequencing did not reveal differences).

In summary, we revealed in this work that a genetically stable industrial yeast strain can exhibit phenotypic variations at least in terms of sugars consumption that likely have a non- genetic origin. Moreover, rare genetic variants (CNVs) can be isolated but they are not detected by standard (not deep) sequencing methods. These CNVs are probably too infrequent to be detected and to produce phenotypic effects but they are still present in the population and could impact more visibly the industrial performance of the strain depending on the environmental conditions. Finally, our work mainly showed that phenotypic variations of non-genetic origin occur, suggesting that the development of specific tools designed to modulate/control such variations, for instance at the gene expression level, are highly relevant in an industrial context. However, it also appears that it might be interesting to develop tools to decrease the basal genetic instability to ensure long-term strain performance.

## Acknowledgements

This work was supported by the Toulouse White Biotechnology (TWB) consortium (pre- competitive financial support NoHighRec to JPC). We thank Tiphaine Clément, Anthony Vollant and Julien Cescut from TWB for their contribution in conducting fermentation experiments on bioreactors; and Alicia Huesca from TBI and Kevin Aere from TWB for their helpful contribution in analytics (HPLC experiments). We also thank Raafat Stephan and Céline Vanwinge (Imaging and flow Cytometry platform of ULiège (GIGA)) for cell sorting performing. Finally, we thank the GeT-Biopuces sequencing platform (Toulouse, France) for performing the quality control of the DNA samples, and the INRAE GENTYANE sequencing platform (Clermont, France), especially Elodie Belmonte for preparing the genomic libraries for long-read sequencing and Vincent Pailler for the bioinformatics analyses.

## Supporting information

Table S1: Evolution of the biomass during the successive batch cultures.

Table S2: List of primers used for qPCR assays.

Table S3: Evolution of the biomass during successive cultivations in microplates. Initial OD was calculated based on the sorted cell number from the estimation that there are about 10^7^ cells/mL for an OD=1.

Table S4: Summary of the genomic variants identified in populations from successive cultivations in microplates. Genotype is indicated as 0 when the variant is absent and as 1 when the variant is present. In brackets is the number of reads getting the variant on the number total of reads at this position. The differences are bolded.

Figure S1: Genome-integrated transgenes of the HDY.GUF12 strain. The different cassettes introduced in the *PYK2* (A), *GAL2* (B) and *HXT2* (C) loci are represented by an arrow, each containing the ORF of the indicated gene and a distinct yeast promoter and terminator. Lines represent the native chromosome. The two distinct chromosomal copies are represented for each locus.

Figure S2: Ethanol tolerance of bulk population from different successive batch cultures. Cells from Duplicate.1 (A) or Duplicate.2 (B) were cultivated in YNB medium containing 20 g/L D-glucose, 10 g/L D-xylose and 10 g/L L-arabinose and supplemented with 31.5 g/L ethanol during 24 h. About 5.10^6^ cells were collected, washed in PBS and stained using fluorescein diacetate (FDA) at a final concentration of 20 µg/mL to assess cell vitality. The number of stained cells was determined by flow cytometry (excitation with the 488 nm blue laser and emission detection with the FL1 533/30 nm filter) using an Accuri C6 Plus cytometer (Becton–Dickinson) (n>=2).

Figure S3: Growth phenotype of clones isolated from successive batch cultures.

A: Principle of the screen applied to identify clones exhibiting a growth defect on xylose and/or arabinose (see material and methods section for details).

B: Spotting assays of serial dilutions of cell suspension on YNB plates containing the indicated sugar at a final concentration of 20g/L.

C: Summary of the number of identified clones per tested batch exhibiting a growth defect on xylose and/or arabinose.

D: Summary of the different growth phenotypes of the clones isolated on plate.

Figure S4: Correlation between C5 sugars consumption and ethanol production of selected clones isolated from batch cultures.

Extracellular xylose, arabinose and ethanol were quantified after 24 h on YNB medium containing 16 g/L D-glucose, 13 g/L D-xylose and 11 g/L L-arabinose (n=2).

Figure S5: Standard curves used for qPCR assays calibration.

Relationship between Cts and the logarithm of the copy number of *ACT1* (A), *araT* (B), *araD* (C), *araB* (D) and *araA* (E). Regression line parameters and percentage of efficiency (E%) based on each experiment are listed on curves.

Figure S6: Gates applied during the cell sorting experiments.

Data from the CO cultivation cycle are shown as an example. Ten thousand events were collected to define the gates applied for subsequent cell sorting.

A: The first gate (P1) was applied to select cells based on YFP (FITC-A) and tdTomato (PE- A) fluorescence signals.

B: The P2 gate was applied on P1-included cells to select single cells with similar cell size and granularity, based on the FSC-A vs. FSC-H plot.

C: Based on the histogram of the fluorescence, single cells with the lowest (P3) and the highest (P4) fluorescence levels were applied and used for cell sorting.

D: Cells number details found in each gate.

Figure S7: Evolution of the fluorescence profile of the strains during the successive cultivations in microplates.

