## Supplementary material for "Long-term phenotypic and genomic instability of an industrial ethanol-producing and C5-utilizing *Saccharomyces cerevisiae* strain": Supp Figures

Figure S1

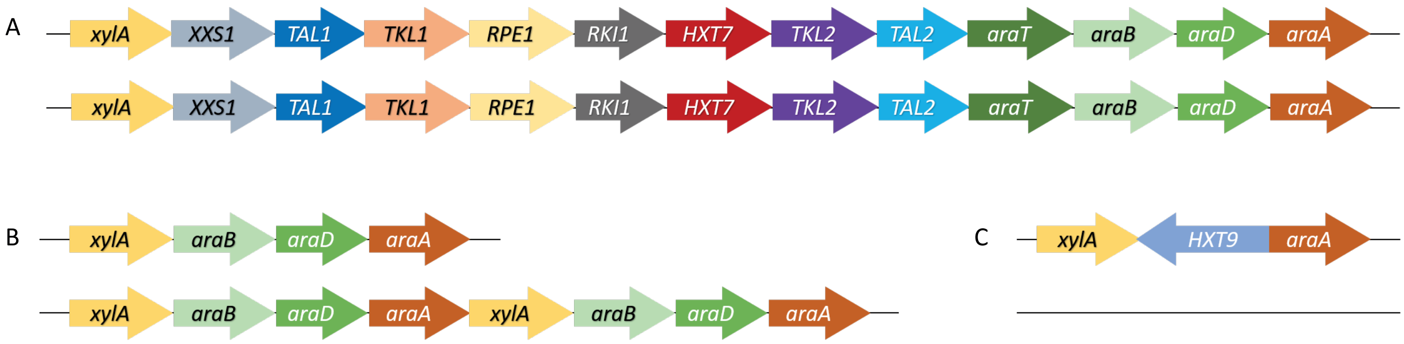

A
xylA XXS1 TAL1 TKL1 RPE1 RKI1 HXT7 TKL2 TAL2 araT araB araD araA
xylA XXS1 TAL1 TKL1 RPE1 RKI1 HXT7 TKL2 TAL2 araT araB araD araA
B
xylA araB araD araA
xylA araB araD araA xylA araB araD araA
C
xylA HXT9 araA

### Figure S2

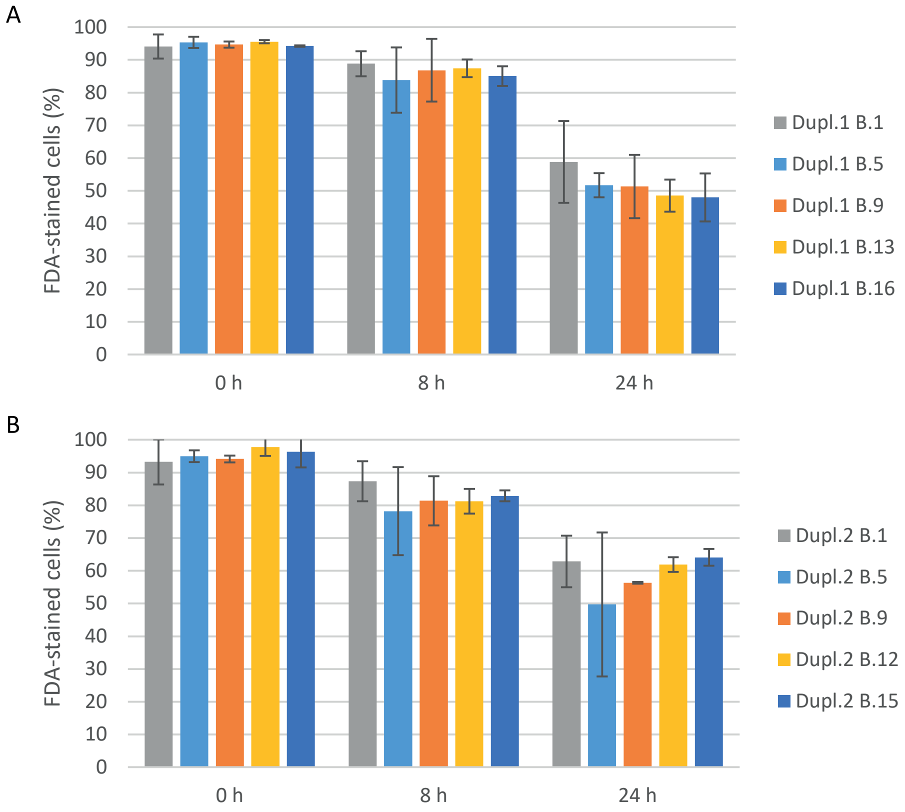

A
FDA-stained cells (%)
100
90
80
70
60
50
40
30
20
10
0
0 h
8 h
24 h
Dupl.1 B.1
Dupl.1 B.5
Dupl.1 B.9
Dupl.1 B.13
Dupl.1 B.16
B
FDA-stained cells (%)
100
90
80
70
60
50
40
30
20
10
0
0 h
8 h
24 h
Dupl.2 B.1
Dupl.2 B.5
Dupl.2 B.9
Dupl.2 B.12
Dupl.2 B.15

### Figure S3

A

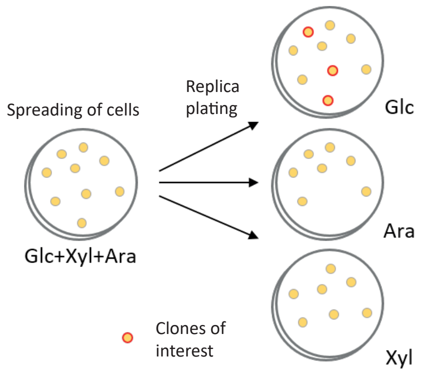


B

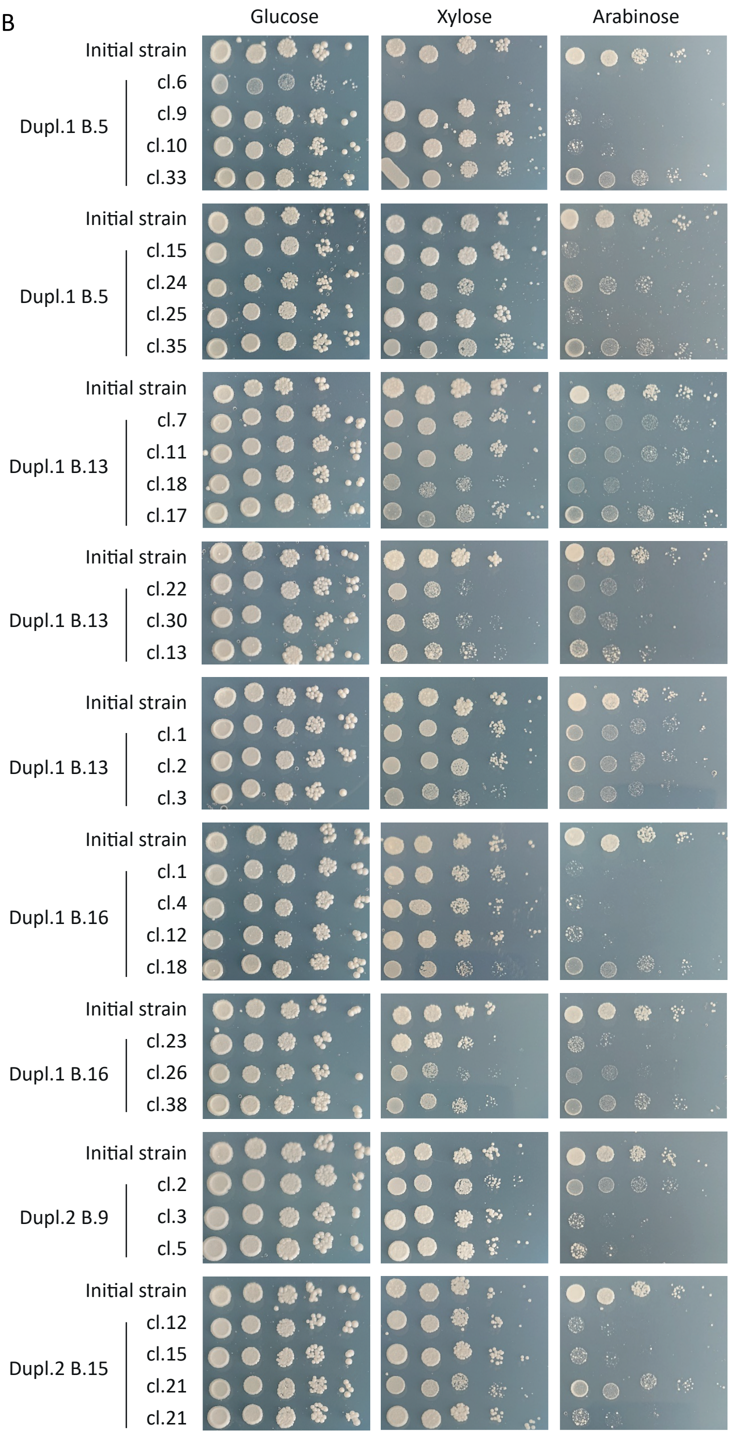


C

| | Batch | Number of isolated clones | % of total |
|---|---|---|---|
| Initial strain | | 0 | 0 |
| Dupl.1 | 5 | 8 | 0,94 |
| | 9 | 0 | 0 |
| | 13 | 10 | 1,49 |
| | 16 | 7 | 0,96 |
| Dupl.2 | 5 | 0 | 0 |
| | 9 | 3 | 0,37 |
| | 12 | 0 | 0 |
| | 15 | 4 | 0,6 |

D

| Number of isolated clones | Intensity of the growth phenotype: Glucose | Xylose | Arabinose |
|---|---|---|---|
| 1 | + | no growth | no growth |
| 10 | wt | wt | +++ |
| 10 | wt | + | + |
| 3 | wt | ++ | + |
| 5 | wt | ++ | +++ |
| 3 | wt | + | +++ |

\+ Low defect
++ Medium defect
+++ High defect

### Figure S4

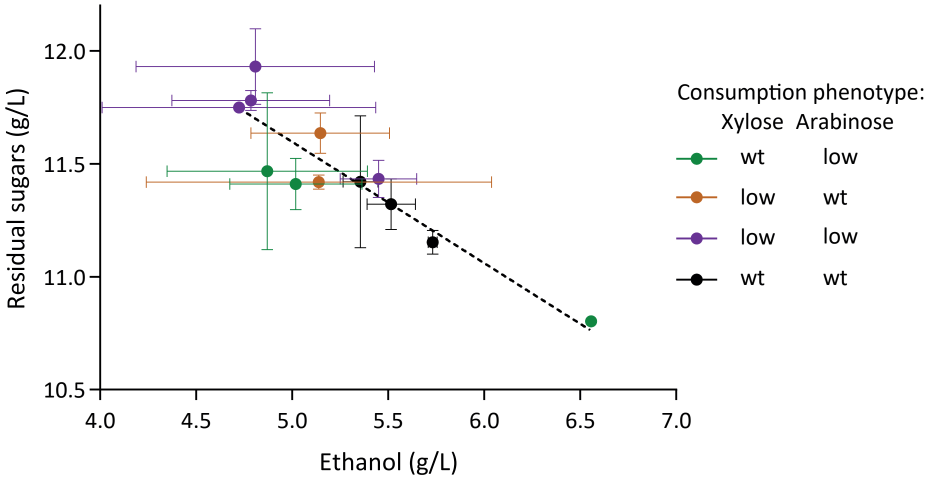

Residual sugars (g/L)
12.0
11.5
11.0
10.5
4.0
4.5
5.0
5.5
6.0
6.5
7.0
Ethanol (g/L)
Consumption phenotype:
Xylose Arabinose
wt low
low wt
low low
wt wt

### Figure S5

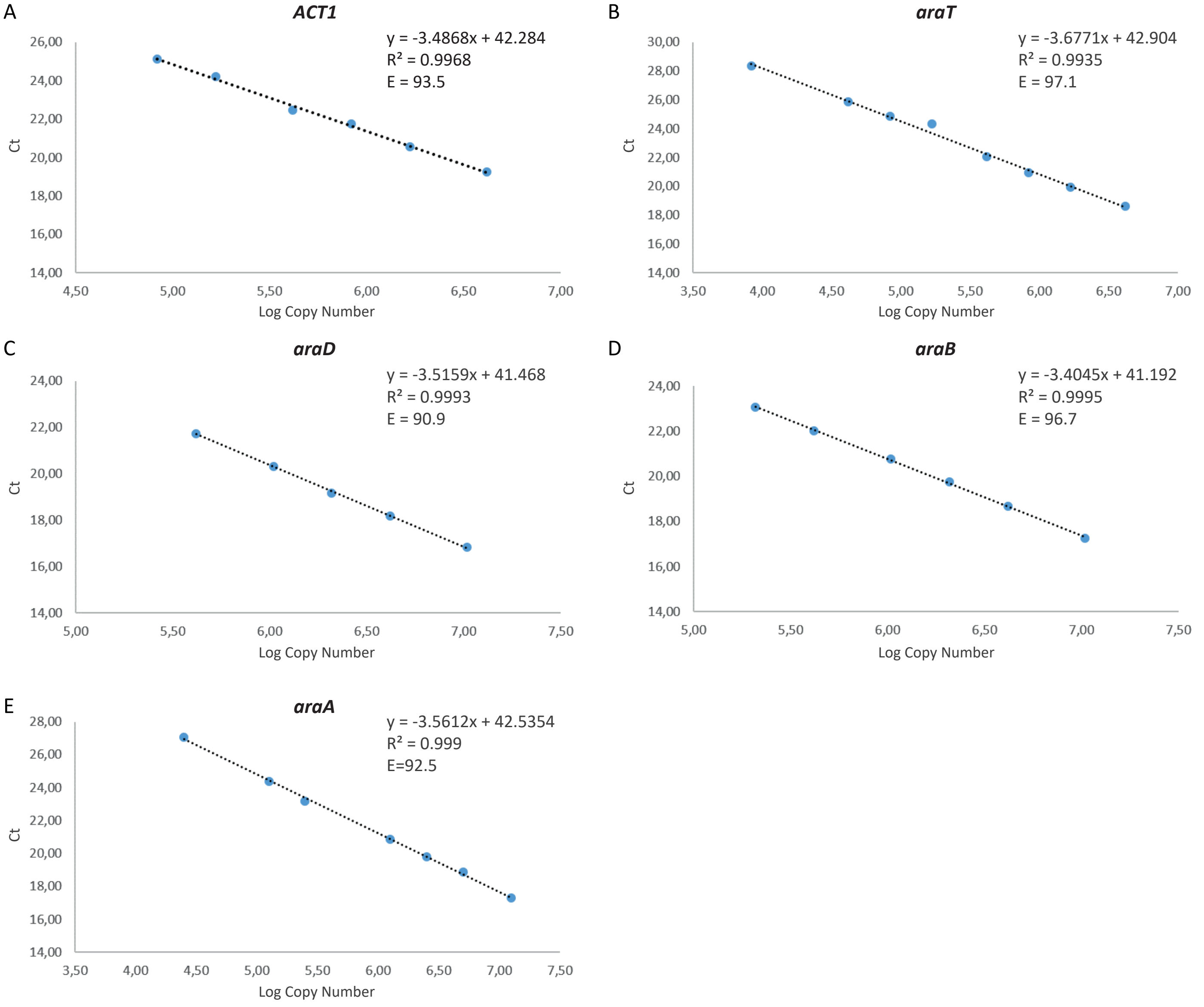
A
ACT1
y = -3.4868x + 42.284
R² = 0.9968
E = 93.5
Ct
Log Copy Number
B
araT
y = -3.6771x + 42.904
R² = 0.9935
E = 97.1
Ct
Log Copy Number
C
araD
y = -3.5159x + 41.468
R² = 0.9993
E = 90.9
Ct
Log Copy Number
D
araB
y = -3.4045x + 41.192
R² = 0.9995
E = 96.7
Ct
Log Copy Number
E
araA
y = -3.5612x + 42.5354
R² = 0.999
E=92.5
Ct
Log Copy Number

### Figure S6

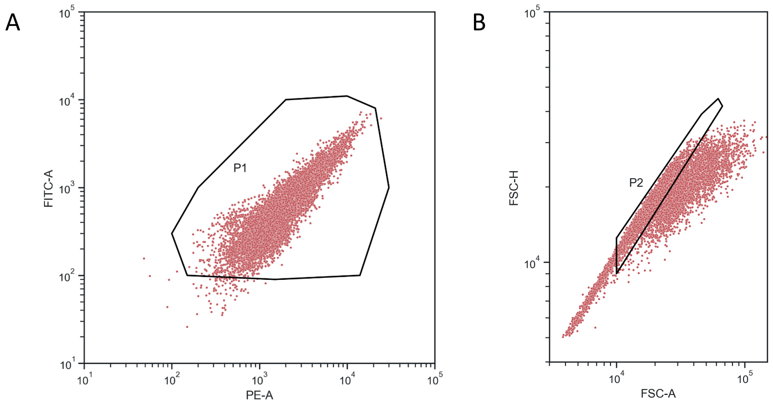


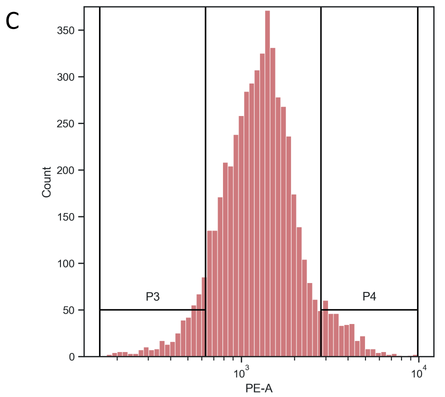


D

| Population | Events | %Parent | %Total | PE-A Mean |
|---|---|---|---|---|
| All events | 10,000 | | 100 | |
| P1 | 9,961 | 99.6 | 99.6 | |
| P2 | 6,292 | 63.2 | 62.9 | 1,395 |
| P3 | 447 | 7.1 | 4.5 | 434 |
| P4 | 402 | 6.4 | 4.0 | 3,490 |

### Figure S7

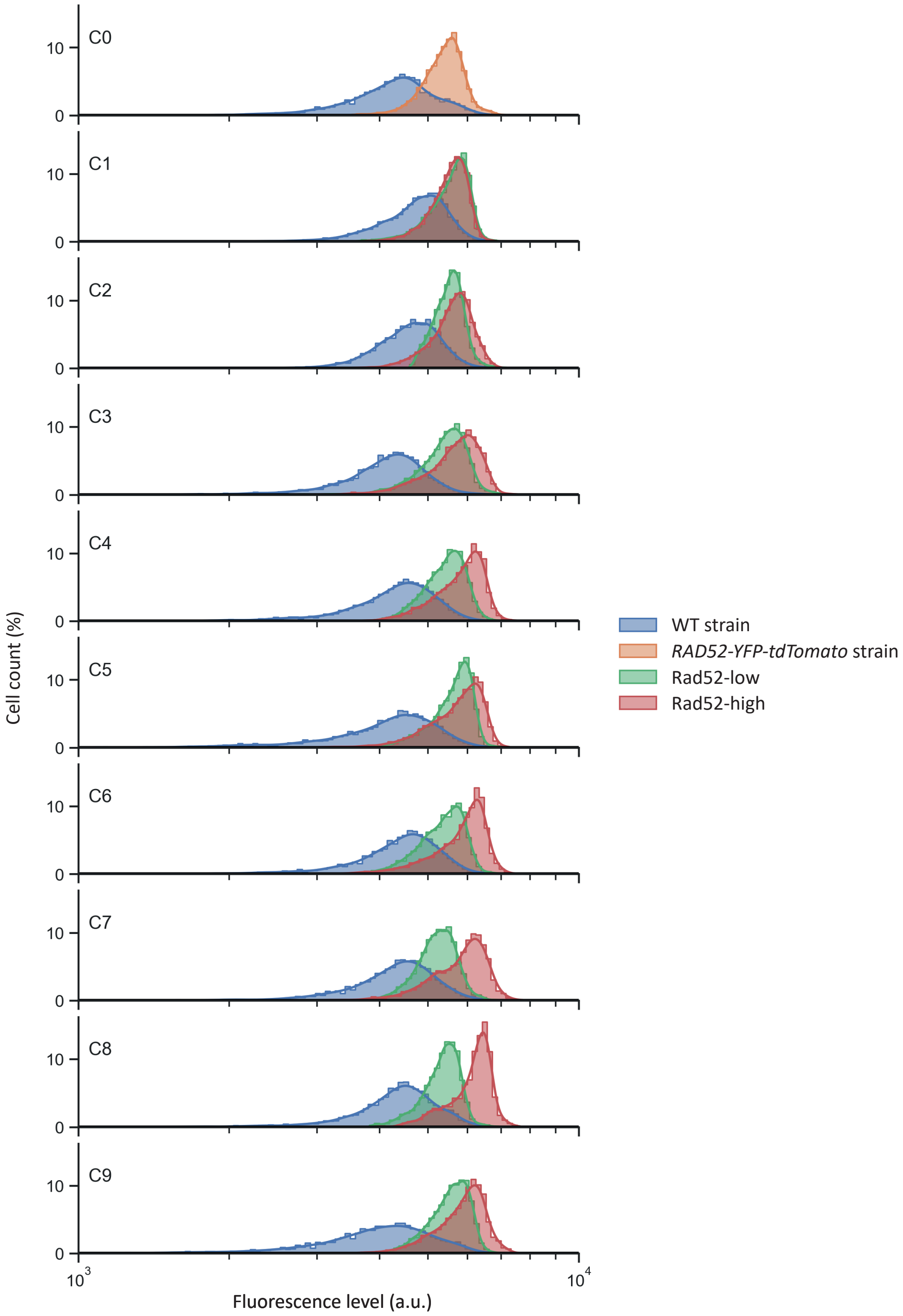
C0
C1
C2
C3
C4
C5
C6
C7
C8
C9
10
0
Cell count (%)
$10^3$
$10^4$
Fluorescence level (a.u.)
WT strain
*RAD52-YFP-tdTomato* strain
Rad52-low
Rad52-high
